# High-throughput iNaturalist image analysis reveals flower color divergence in *Monarda fistulosa*

**DOI:** 10.1101/2025.05.21.655392

**Authors:** Patrick F. McKenzie, Samuel H. Church, Robin Hopkins

**Affiliations:** Department of Organismic and Evolutionary Biology, The Arnold Arboretum, Harvard University, Cambridge, MA 02138, USA; Department of Ecology and Evolutionary Biology, Yale University, New Haven, CT, 06511, USA

**Keywords:** Monarda, iNaturalist, flower color, community science, computer vision

## Abstract

Characterizing patterns of trait variation across widespread species is a fundamental goal of natural history. Here we create a pipeline to analyze a large community science dataset and test hypothesized flower color divergence across the range of a widespread wildflower. *Monarda fistulosa* is a North American perennial that produces showy lavender inflorescences. Although previous literature suggests that the flowers of western *M. fistulosa* might display a deeper purple color than the eastern varieties, this divergence has not been assessed at scale. We process over 40,000 community science photographs of *M. fistulosa* to identify flowers and extract color. We demonstrate that the flowers of the montane western variety have lower lightness and higher chroma, corresponding to a deeper violet color, than those of eastern *M. fistulosa*. Our approach and validation provides a scalable framework for phenotyping community science images and enables analysis of geographic color variation in other widespread species.

## Introduction

A foundational goal of natural history has been characterizing patterns in biological diversity. We are entering a new era of natural history, shaped by community scientists documenting biodiversity and by expanding computational tools for large-scale data analysis (Tosa et al. 2021; Seltzer 2019; Aristeidou et al. 2021; Di Cecco et al. 2021). Together, these advances present new opportunities for studying diversity at scale. We leverage these tools to quantify systematic phenotypic variation across the range of the widespread, ecologically and economically important flowering plant species *Monarda fistulosa* L. (Lamiaceae).

*M. fistulosa* is valued in horticulture, native habitat restoration, and traditional food and medicine (Davidson 2007; Quinlan et al. 2021). Its lavender inflorescences attract diverse insect communities, including bees, butterflies, syrphid flies, wasps, beetles, true bugs, and spiders (Keefover-Ring 2022; Smith 2023; Cruden, Hermanutz, and Shuttleworth 1984). Like many herbaceous perennials of North America, *M. fistulosa* is widespread across broad environmental niches and comprises an important carpet of herbaceous biomass throughout the spring, summer, and fall.

While *M. fistulosa* has a broad geographic range and extensive variability, it displays general patterns that have been used to delimit three varieties: two in the eastern U.S. (var. *fistulosa* and var. *mollis* (L.) Benth.) and one in the western U.S (var. *menthifolia* (Graham) Fernald). The eastern varieties are separated by leaf texture and pubescence and are notoriously challenging to distinguish reliably, so the taxonomic integrity of these varieties remains debated (McClintock and Epling 1942; Scora 1967; Turner 1994). In contrast, the western var. *menthifolia* is more distinct, characterized by vegetative traits such as single, usually unbranched stems and shorter leaf petioles (Scora 1967). This variety has maintained stable taxonomic recognition since Fernald (1944).

Vegetative differences might not be the only distinction between the eastern and western *M. fistulosa* varieties. Anecdotal observations suggest a possible difference in flower color, with western populations displaying deeper purple flowers compared to the paler lavender corollas of eastern plants. This difference has received little attention, mentioned only briefly by Wooton (1898), who noted darker flower color in his description of *M. stricta* (described from New Mexico, now synonymous with *M. fistulosa* var. *menthifolia*) relative to the eastern *M. scabra* (described from Missouri, now synonymous with *M. fistulosa* var. *fistulosa*), and by Turner (1994), who differentiated the varieties in his treatment for *Monarda* of Texas in part using flower color descriptors (“corollas deep pink to lavender” for *M. fistulosa* var. *menthifolia* vs “corollas pale pink to pale lavender”). These descriptions suggest subtle, continuous variation in flower color, but this putative west–east divergence has yet to be rigorously evaluated at scale.

Even subtle flower color variation can be detectable by pollinators, shaped by selection, and/or reflective of broader patterns of divergence (Trunschke et al. 2021; Rausher 2008; Sapir, Gallagher, and Senden 2021). Floral traits often play crucial roles in reproductive isolation and speciation; consistent flower color differences between varieties could therefore indicate divergence in reproductive strategies and pollinator interactions, or they could instead reflect unrelated patterns of genetic isolation and drift (Rausher 2008; White and Royer 2024).

Understanding the geographic structure of phenotypic variation is an essential first step in addressing these questions. Studying flower color typically requires performing pigment extractions or standardized photography. These approaches require time, financial resources, and access to private and restricted public lands. However, community science platforms like iNaturalist now aggregate millions of georeferenced images of flowering plants worldwide, creating new opportunities to address certain questions involving color (iNaturalist.org, n.d.; Laitly et al. 2021). Meanwhile, advances in computational image analysis enable efficient, large-scale phenotyping from these datasets (McKenzie, Berardi, and Hopkins 2025).

To test for flower color divergence between regions, we analyzed over 40,000 *M. fistulosa* images from iNaturalist. We created a pipeline combining general-purpose computer vision with image segmentation for high-throughput phenotyping, and we rigorously validated its performance. This enabled us to map spatial color patterns and confirm deeper purple corollas in western *M. fistulosa*.

## Methods

### Data Acquisition

#### GBIF export of iNaturalist observations

Community scientists have documented thousands of observations of *M. fistulosa* through iNaturalist, and the research-grade and licensed subset of observations, metadata, and photographs are consolidated on GBIF, a large biodiversity database (GBIF.org, n.d.). We exported all research-grade *M. fistulosa* observations from GBIF, yielding tables corresponding to 1) all observations, and 2) all corresponding images. Each image maps to a specific observation, while each observation could potentially map to multiple images.

#### Image download and mapping

We downloaded all images associated with these observations using a python script that individually queried each URL in the GBIF table of images. We retained mappings between each image and its corresponding observation’s metadata containing date, latitude, and longitude.

### Image preprocessing

#### Image classification (GPT-4o)

We used a general purpose computer vision model to filter downloaded images to those containing *Monarda* (common name “beebalm”) flowers. Specifically, using a python script we submitted each photo to the OpenAI API’s GPT-4o model to classify each image (OpenAI et al. 2024). The query used was “Answer YES or NO: Is this a high-quality close-up photo of a beebalm flower?” We then filtered the dataset to those images for which the model returned the answer “YES.”

#### Semantic segmentation (Roboflow)

We extracted flowers from each photo in the filtered dataset using a trained semantic segmentation model. Semantic segmentation models classify every pixel in an image as belonging to one of a distinct number of categories. For our model, we specified that each pixel should take one of two categories: “flower” or “not a flower.” We first used the built-in annotation tools, driven by Segment Anything (SAM), on the platform Roboflow to manually label a subset of 110 pictures from our flower dataset (Roboflow, n.d.; Meta AI, n.d.). This annotation process involved clicking on the images inside the Roboflow annotation GUI: a single click on a flower created a polygon that approximated the boundary of that feature; to the extent that the polygon encompassed too broad or too restricted of an area, we clicked in the area that should be excluded or included to adjust the polygon shape. We were conservative while annotating, labeling the obvious flowers but tending toward exclusion in edge cases in which there was uncertainty about whether pixels represented flowers. Our intention was to maximize the model’s true-positive rate, since our ultimate goal was to sample *most* flower pixels, without necessarily needing to sample *all* flower pixels. We split our annotated dataset into 77 images for model training, 22 for validation, and 11 for testing. We applied the automated training method for the Roboflow 2.0 Semantic Segmentation model, with default parameters and the only preprocessing step being the recommended “auto-orientation” option. The trained model accepts an input image and returns a category for each pixel in that image.

We used a python script to query the trained model, which was hosted on the Roboflow online platform, for each image in the GPT-filtered dataset. We saved each resulting json mask (holding the model’s pixel labels for a particular image) into a local directory. Finally, in python, we extracted the pixel color values of the designated “flower” pixels from each image.

### Color phenotype assignment

For the extracted flower pixels from each image, we calculated the geometric median (robust to outliers) of the CIELAB components (a three-dimensional color space approximating perceptual uniformity, therefore facilitating numerical operations) of the extracted pixels using an efficient algorithm (Vardi and Zhang 2000). We converted this color to its associated LCh value (representing the same color space, but in polar coordinates) value for downstream analyses and visualizations using the python packages opencv-python and python-colormath (OpenCV, n.d.; Taylor, n.d.). Through our analyses, we used CIELAB coordinates for numerical operations and LCh values for visual interpretation, since its components (lightness, chroma, and hue) are intuitive (Boronkay et al. 2024).

### Spatial aggregation and west vs. east analysis

For any iNaturalist observation having multiple images of flowers, we randomly sampled one image to use for downstream analysis. We used the latitude and longitude values associated with the iNaturalist observation for mapping.

To visualize spatial trends in flower color, we directly mapped observations using the CIELAB color (converted to hex code) inferred for that flower to represent the observation. To simplify visualization, we also averaged the CIELAB values of observations within 200km x 200km grid cells to summarize the dominant color of flowers in each location.

Finally, we used Hotelling’s T^2^, a multivariate extension of the t-test, to compare color components on either side of a -100° longitude breakpoint roughly separating *M. fistulosa* var. *fistulosa* from *M. fistulosa* var. *menthifolia*, testing the hypothesis that the west and east do not differ in color (Hotelling 1992). We visualized the differences between these regions using a transformation into LCh color space, allowing us to look at variation in hue, chroma, and lightness.

### Validation analysis

#### Segmentation accuracy

To validate our results, we first confirmed that our segmentation protocol accurately identified flower petals. All *M. fistulosa* flowers are shades of purple; we therefore assessed the distribution of assigned phenotypes in CIELAB space and in LCh space to determine if they are, in fact, all shades of purple (ranging from rare white individuals, to common light lavender and deeper violet).

#### *Recovering the* M. fistulosa *var.* menthifolia *boundary*

Next, we evaluated whether the detected color divergence emerges independently of any predefined boundary between *M. fistulosa* varieties. First, we performed sliding Hotelling’s T^2^ tests to find the longitude value that yields the best explanatory power (i.e. the highest test value) when separating the data into western vs. eastern regions. Specifically, we tested 1000 values evenly spaced between the minimum and maximum longitudes in our dataset (−135.3° and - 61.4°), requiring a minimum of 100 observations in each population to perform the test. We used the three CIELAB components for each observation as our response variables, and we identified the longitude that maximized Hotelling’s T^2^ as implemented in the python package pingouin (Vallat 2018).

Second, we used k-means to naively cluster the full dataset into k=2 categories using only the CIELAB color for each observation. One cluster was assigned a value of 1, and the other was assigned a value of 0. We then extracted windows centered at each of the same 1000 evenly spaced values, with each window including all observations +/-1 degree of the focal longitude, and we averaged the assigned cluster values of points in each window to identify shifts in proportion of observations assigned to cluster 1 versus cluster 0.

#### Lighting bias assessment

Finally, we considered whether color differences between western and eastern images could be due to systematic differences in lighting conditions between the regions. We hypothesized that sunnier conditions could lighten the perception of flower color due to overexposure in the photograph. To assess this, we created a validation subset of images by randomly selecting 500 images from the primary dataset: 250 from the west (<-100° longitude) and 250 from the east (>-100° longitude). Two authors with little prior knowledge of the species or sampling strategy (S.C. and R.H.) independently binned each image into one of three categories to indicate the lighting of the flowers: shade, partial sun, or full sun. The authors were also permitted to answer “NAN” if they felt unable to categorize a particular image. Finally, to assess the accuracy of the iNaturalist data and GPT labeling, each author was instructed to note whether any of the photos did not feature a *Monarda* flower.

We analyzed our binning results using two-way ANOVAs and a joint MANOVA to test for differences in color due to main effects of lighting condition and region (west vs. east) and for the interaction between lighting condition and region. A nonsignificant interaction effect indicates there is no systematic difference in the effect of region on color across the different lighting conditions.

## Results

### Data Acquisition

The GBIF export yielded 28,750 research-grade observations of *M. fistulosa* with 41,069 corresponding images (19 February 2025: https://doi.org/10.15468/dl.v64sz8).

### Image preprocesing

#### Image classification

Of the 41,069 images in the dataset, GPT classification identified 20,761 as being high-quality pictures of flowers (Figure 1A-B) [notebooks_pipeline notebooks 1, 2, 3]. We assessed a 500-image random subset of these images and determined that all 500 contained *Monarda* flowers.

**Figure 1:**
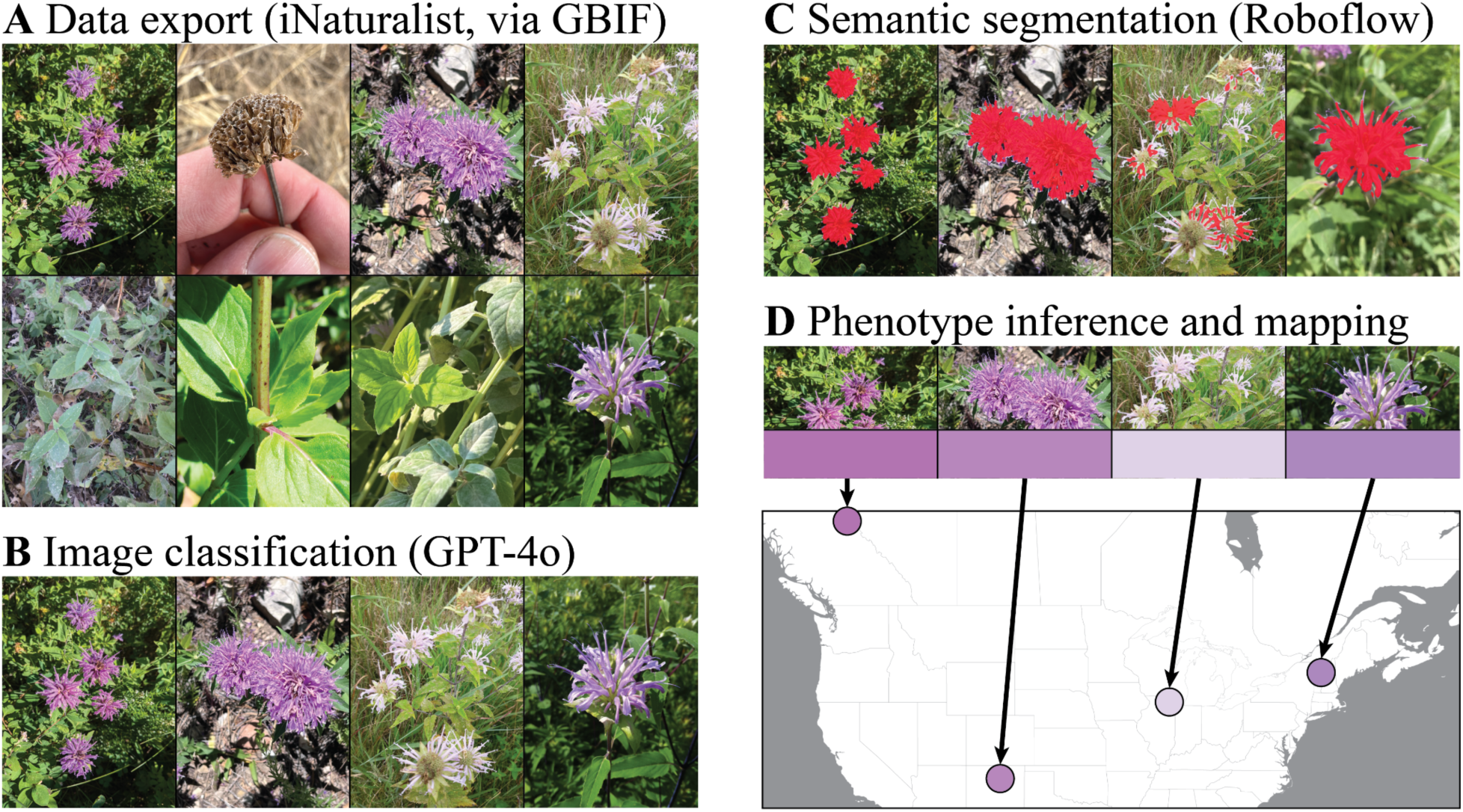
Steps with representative photographs of the flower color phenotyping pipeline. A) We exported all research-grade iNaturalist observations of *Monarda fistulosa* from GBIF. B) We used GPT-4o to classify each image as to whether it contained a flower. C) We trained a Roboflow semantic segmentation model on a subset of images and applied the trained model to extract “flower” pixels from each image in the dataset. D) We calculated the geometric median of each set of extracted pixels to represent the flower color phenotype from each image, and we paired this phenotype with the observation’s iNaturalist metadata for spatial analysis.

#### Semantic segmentation

The annotation and training of the segmentation model required <5 hours of manual labeling, and the automated model trained in under 30 minutes. The model trained to 82.5% MIoU (Mean Intersection-over-Union), indicating the degree to which the predicted pixels containing flowers overlapped with those that we had manually annotated for images in the test dataset. The trained model yielded a pool of pixels for every image that was “enriched” for the flower color, from which we inferred a single phenotype (Figure 1C-D). The total dataset contained 20,628 images with extracted pixels and corresponding colors (133 failed due to not detecting flowers or from file format errors). These images mapped to 16,587 unique georeferenced iNaturalist observations [notebooks_pipeline notebooks 4, 5, 6].

### Spatial aggregation and west vs. east analysis

We mapped the color for each image to the corresponding iNaturalist observation, binned the observations in 200km x 200km blocks, and calculated an average CIELAB value for each block. The resulting map highlights color differences between the west, which features deeper violet colors, and the east, whose assigned phenotypes are consistently more pale lavender (Figure 2A, individual observations shown in Figure S1).

**Figure 2:**
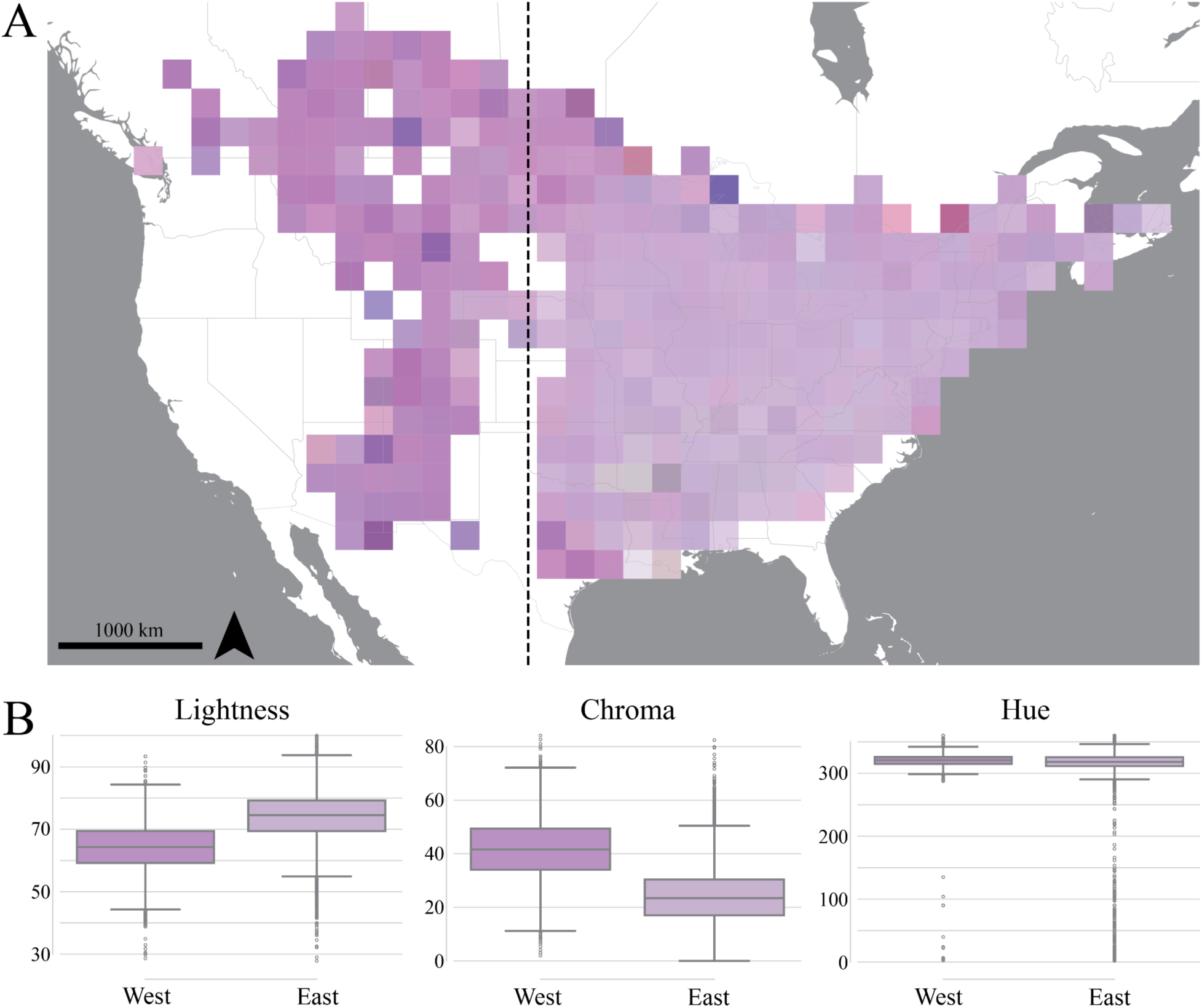
Spatial summary of *Monarda fistulosa* flower colors across North America. A) Map of the color of *M. fistulosa* flowers. Each square is a 200km x 200km cell with the color corresponding to the average median CIELAB color value of each observation in the cell. The dotted line denotes -100° longitude, separating eastern and western regions of the range. B) Boxplots summarizing LCh color components west and east of -100° longitude, with each box showing the median and interquartile range, and with the color of each box reflecting the geometric median CIELAB value from west and east, respectively.

There is a significant and large effect of region (i.e. west vs. east) on color as revealed by a two-sample Hotelling’s T^2^ test comparing the distributions of the three LCh and CIELAB color components on either side of -100° longitude (CIELAB: T^2^=6,368.12, F (3, 16,583)=2,122.45, p < 0.001, partial-eta^2^=27.7%; LCh: T^2^=5978.90, F (3, 16,583)=1992.73, p < 0.001, partial-eta^2^=26.5%) [notebooks_figures/figure_2b.ipynb]. We visualized differences between the two regions for the three LCh components (Figure 2B) and for each CIELAB component individually (Figure S2). Notably, comparing flower color between west vs. east in LCh space showed that the *hue* (i.e., “h”) component remains similar between regions, while lightness and chroma exhibit stark differences.

### Validation analysis

#### Segmentation accuracy

We validated that the segmentation model successfully identified flowers by confirming that nearly all observations fell in purple color space (Figure 3A). This is plotted in a three-dimensional histogram with LCh choma (C) and hue (h) on the x- and y-axes and with observation frequency on the z-axis. In LCh color space, shades of purple and violet correspond to *h* values (hue) in 0-50 and 250-360. As expected, nearly all observations fell within this space, with only 119 observations (0.7% of overall) in the range *h*=[50,250] (Figure 3B) [notebooks_figures/figure_3b.ipynb]. These outliers appeared at high lightness and low chroma (Figure S4). Visual inspection of the outliers reveals true biological variation, actually identifying aberrant white *M. fistulosa* individuals that have previously been suggested at varietal status (var. *albescens*), and for which the inferred hue is highly sensitive to lighting conditions (Farwell 1923).

**Figure 3:**
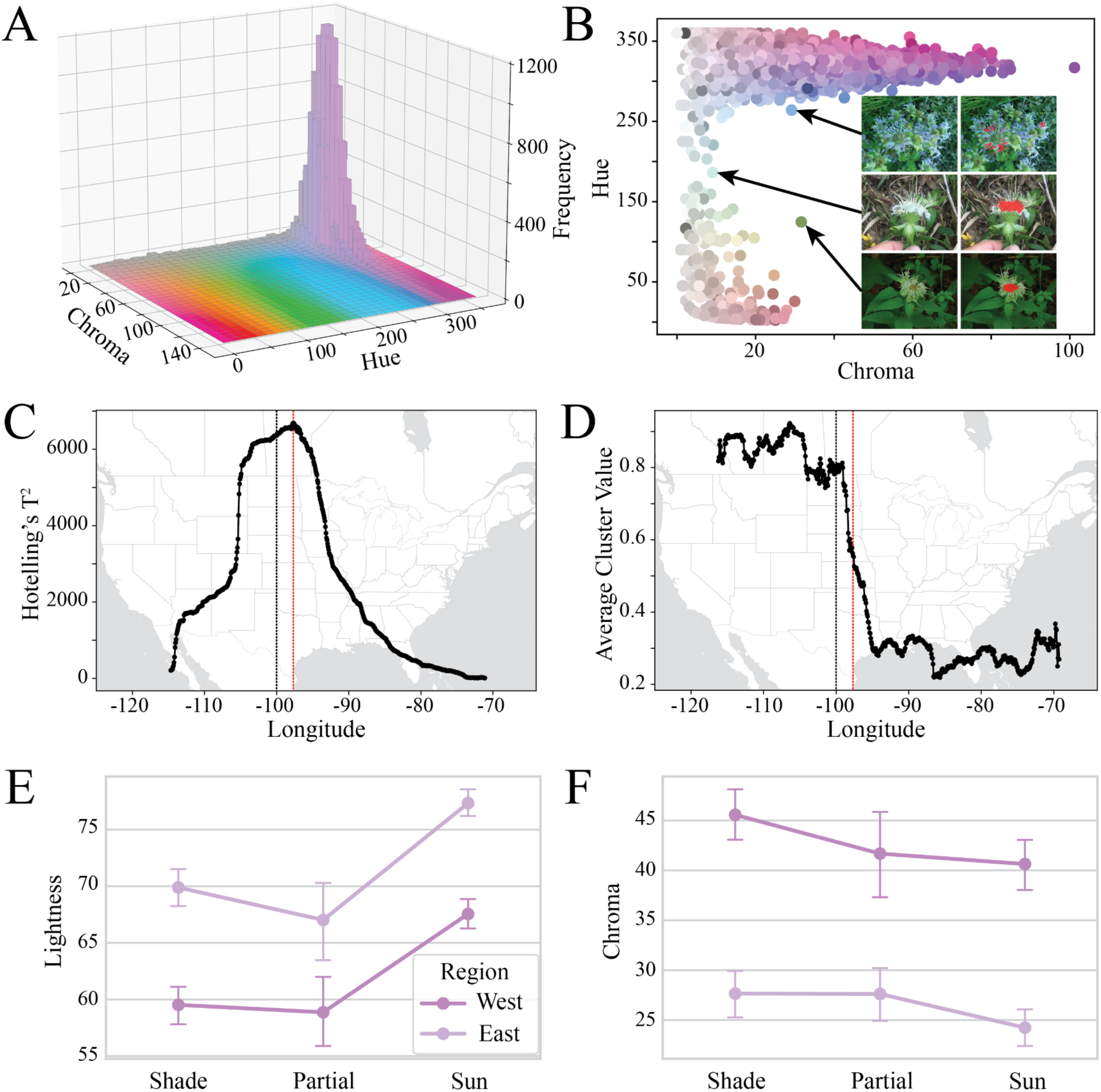
Results from validation analysis. A) Three-dimensional histogram of *M. fistulosa* flower colors in our dataset showing frequency of observations across values of chroma and hue from LCh color space (lightness held constant at value 70). Complete rotation shown in Figure S3. B) Two-dimensional projection of LCh hue vs. chroma, highlighting outlier points falling along the hue axis in the range h=[50-250]. Inset displays representative photographs of outlier points and corresponding segmentation demonstrating that outliers are typically real biological variation or lighting variation, rather than segmentation errors. See Figure S4 for all pairwise two-dimensional representations of LCh color space. C) Sliding Hotelling’s T^2^ statistics on CIELAB color components of *M. fistulosa* observations split into west vs. east regions across 1,000 longitudinal divides between -135.3° and -61.4°. The red dotted line indicates the peak Hotelling’s T^2^ value at longitude of -97.67°, and the black dotted line indicates -100° longitude. D) Sliding window average of cluster assignment from k-means clustering on CIELAB color components with k=2. Each window spanned two degrees longitude, and the red dotted line and black dotted line are stationary from the previous panel. The dramatic drop in value reveals a shift in cluster assignment from 1 to 0, moving west to east, between longitudes of -99° and -95°. E-F) Point plots showing how the mean lightness (E) and chroma (F) vary across lighting categories and geographic regions. Each point represents the mean value for its group, with error bars indicating 95% confidence intervals.

#### *Recovering the* M. fistulosa *var.* menthifolia *boundary*

We used two approaches to naively subdivide the population as though we had no prior knowledge of the west/east distribution of *M. fistulosa* varieties. Using a sliding multivariate t-test (Hotelling’s T^2^), we found a clear peak at longitude = -97.67° (Figure 3C). Breaking the data into two populations using this longitude explained 28.7% of the multivariate variance (partial eta-squared) in CIELAB color components [notebooks_figures/figure_3c.ipynb].

Using k-means clustering with k=2 on the CIELAB color components across all observations, it was visually apparent that the data clustered into western and eastern populations (Figure S5). Sliding window averaging of cluster assignments found that a high proportion (>80%) fell into cluster 1 in the west, with a sharp transition down to cluster 0 in the east (<30%) occurring from -99° to -95° longitude (Figure 3D).

#### Lighting bias assessment

We found that the biological signal of relative differences in flower color is robust to the noise inherent in community science photographs. Two authors showed 74.2% agreement in binning 500 images into three lighting categories: sun, shade, and partial sun (Cohen’s k = 0.586, 95% CI 0.525-0.645) [notebooks_figures/figure_3ef.ipynb]. In the consensus dataset of agreed-upon lighting category (n=371), lighting conditions were not significantly different between the west and east (p=0.78, chi-square=0.51) [notebooks_figures/figure_3ef.ipynb].

Formal model testing of differences in CIELAB color components with a joint MANOVA revealed that the combination of flower color components differed significantly by region (Pillai’s trace = 0.082, F(3, 363) = 10.85, p < 0.001) and by lighting category (Pillai’s trace = 0.322, F(6, 728) = 23.28, p < 0.001), but the interaction between region and lighting was not significant. In univariate ANOVAs, region explained 25–41% of the variance in each CIELAB component, whereas lighting explained 1–27% (partial-eta-squared). Lighting category had its strongest individual effect on color lightness (L). As expected, relative to flowers photographed in partial sun and shade, colors inferred from full-sun flowers were much lighter (β = 10.32 units in L, p<0.001).The interaction between region and lighting was not significant for any test, indicating that the west-east difference is consistent across lighting categories [notebooks_figures/figure_3ef.ipynb].

## Discussion

We find that western, montane *M. fistulosa* var. *menthifolia* produce deeper violet corollas than eastern *M. fistulosa*. This pattern is driven by clear differences in lightness and chroma between regions. We create an efficient, reproducible pipeline to quantitatively assess flower color across the range of *M. fistulosa* from over 40,000 community science images, and we confirm the robustness and potential breadth of application of our method by performing a series of validation analyses.

Using our phenotyping pipeline, we discovered a systematic difference in flower color marked by a known varietal boundary. This phenotype transition provides a valuable reference for delimiting boundaries between *M. fistulosa* var. *fistulosa* and *M. fistulosa* var. *menthifolia*, supplementing previous work using herbarium specimens (Scora 1967). Our findings motivate future work around this transitional zone to examine genetic divergence and admixture between varieties, potentially correlating flower color clines with genomic patterns of differentiation and with clines in other characters.

Biologically, the divergence in flower color between west and east presents some interesting hypotheses. The darker pigmentation in western plants could reflect a response to their higher elevation habitats. Previous research in other systems suggests that anthocyanin pigment production can be protective against the high ultraviolet radiation exposure of mountain habitats (although typically in foliar tissue) (Berardi et al. 2016; Chalker-Scott 1999). Further studies could leverage our pipeline to examine flower color patterns in other wide-ranging, anthocyanin-producing species to test the generality of this pattern. Additionally, as evidenced by extensive research in other systems (Milano, Kenney, and Juenger 2016; Barazani et al. 2019; Lau and Galloway 2004), the evolution of flower color can have profound impacts on pollinator interactions, changing both the species that visit flowers and the foraging behavior of visitors (Sapir, Gallagher, and Senden 2021; Willmer 2011; Rausher 2008). Pollinator behavioral response to floral variation has important implications for patterns of gene flow and reproduction both within and between species (Burgin and Hopkins 2022; Wessinger 2021; Hopkins 2022). However, relatively little research has studied the impact of continuous (vs. discrete) color variation on pollinator interactions (Trunschke et al. 2021; Sapir, Gallagher, and Senden 2021). Our pipeline thus provides a robust means of quantifying continuous variation in color and could enable future research characterizing the evolutionary forces driving this variation and its ecological consequences.

Our results highlight the value of community science platforms for large-scale phenotypic research. While there is variability inherent in user-submitted photographs, we were most interested in the *relative* shade of flowers between regions, and this pattern emerged through the noise. Our series of validation analyses supported our findings as reflecting true biological signal rather than technical artifacts. Our pipeline balanced the specificity required to sample *Monarda* flower color with scalability through the use of models that required minimal supervision. Similar pipelines have been developed for extracting color from community science data and might have reached the same conclusions, but for this analysis existing pipelines seemed either overly time-intensive, such as as requiring manual clicking of images (Luong et al. 2023), or overly general and lacking specificity on *Monarda* flowers (Boyko 2024; Perez-Udell, Udell, and Chang 2023). Finally, while we reduced the flower color in each image to one value, future approaches might instead analyze the flower color for each image as a distribution of values.

Our analysis serves as a demonstration of how advancements in technology and community science have broadened opportunities in natural history research, enabling analyses previously impractical due to logistical constraints. The extensive digital databases of georeferenced plant images available today are unprecedented resources, and probing them with emerging computational tools will allow botanists and ecologists to explore patterns of biodiversity and trait evolution at unprecedented scales.

## Acknowledgements

The iNaturalist and GBIF platforms enabled this analysis. We are especially grateful to the community of iNaturalist users, curators, and staff who generated and maintain the data. R.H. was funded by NIH NIGMS (1R35GM142742-01). P.F.M. and R.H. were funded by the Harvard University Dean’s Competitive Fund and the OpenAI Researcher Access Program.

## Statement of Authorship

P.F.M. conceived of the project. P.F.M., S.H.C., and R.H. designed the methods. P.F.M., S.H.C., and R.H. performed the analysis. P.F.M. wrote the original manuscript. P.F.M., S.H.C, and R.H. reviewed and edited the manuscript.

## Data and Code Availability

All code required for the analysis (referenced with brackets in text above), as well as datasets produced by the analysis, are available at: https://github.com/pmckenz1/monarda_fistulosa_color The trained Roboflow segmentation model is available at: https://universe.roboflow.com/patricks-dashboard/monarda_fistulosa_segmentation/model/1

---

**Figure S1:**
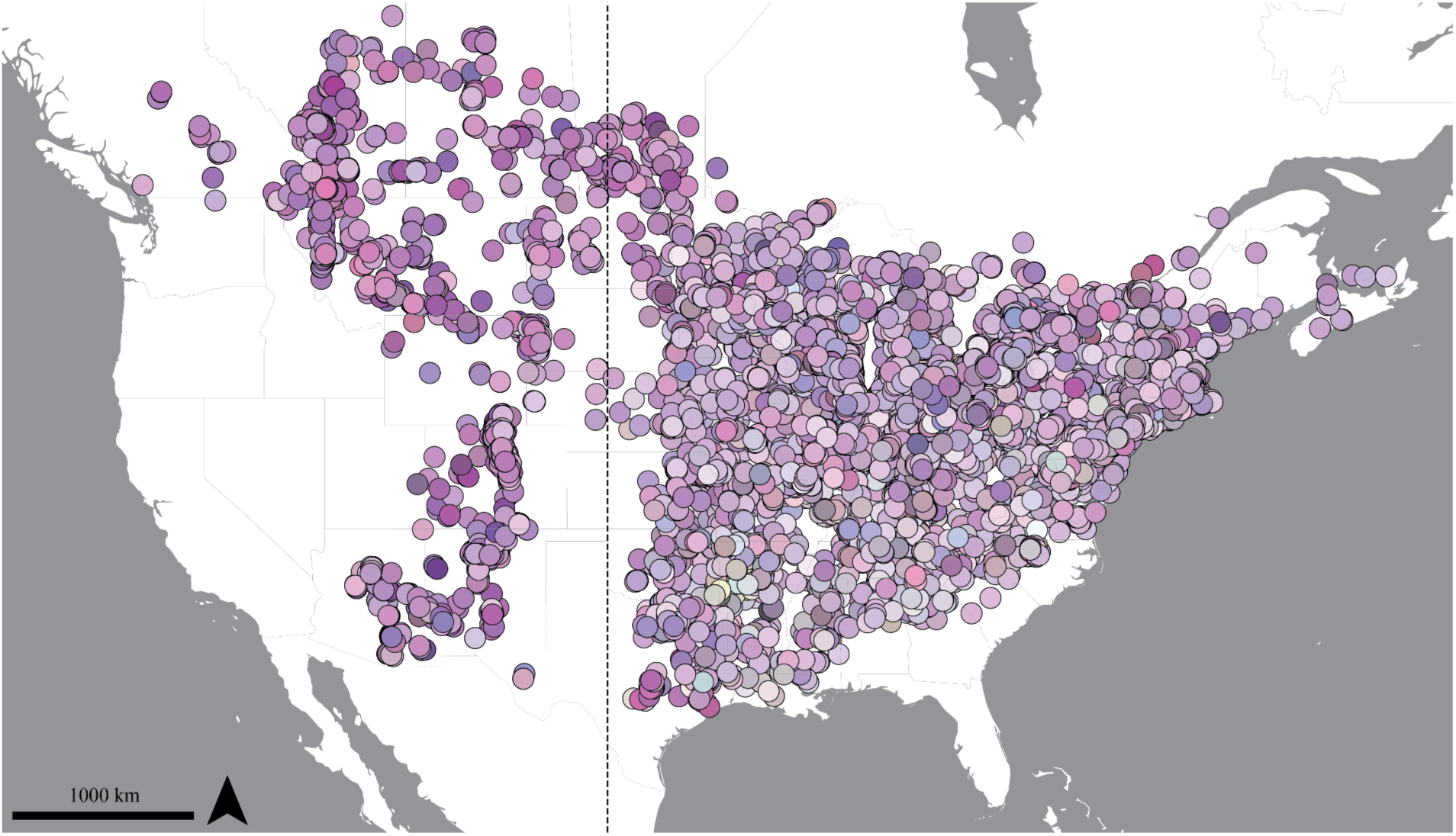
All individual iNaturalist observations mapped using their inferred color. These individual observations underlie the grid-averaged result in Figure 2A. The dotted line indicates - 100° longitude.

**Figure S2:**
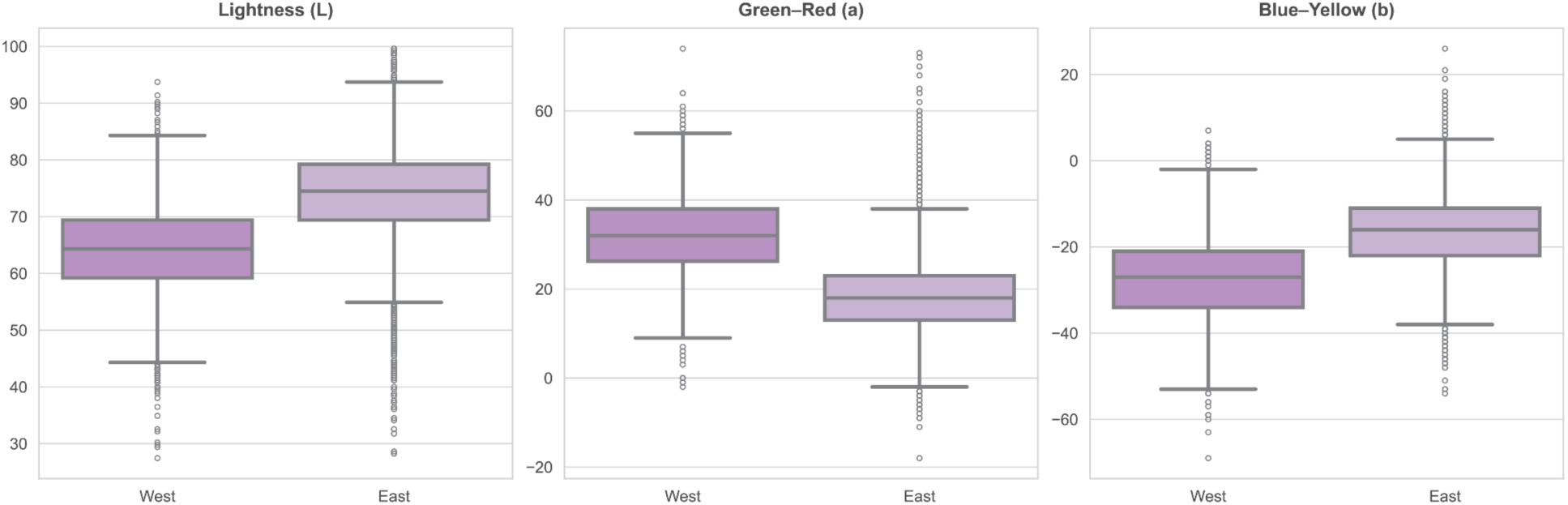
Boxplots summarizing CIELAB color components west and east of -100° longitude, with each box showing the median and interquartile range, and with the color of each box reflecting the geometric median CIELAB value from west and east, respectively.

**Figure S3:**
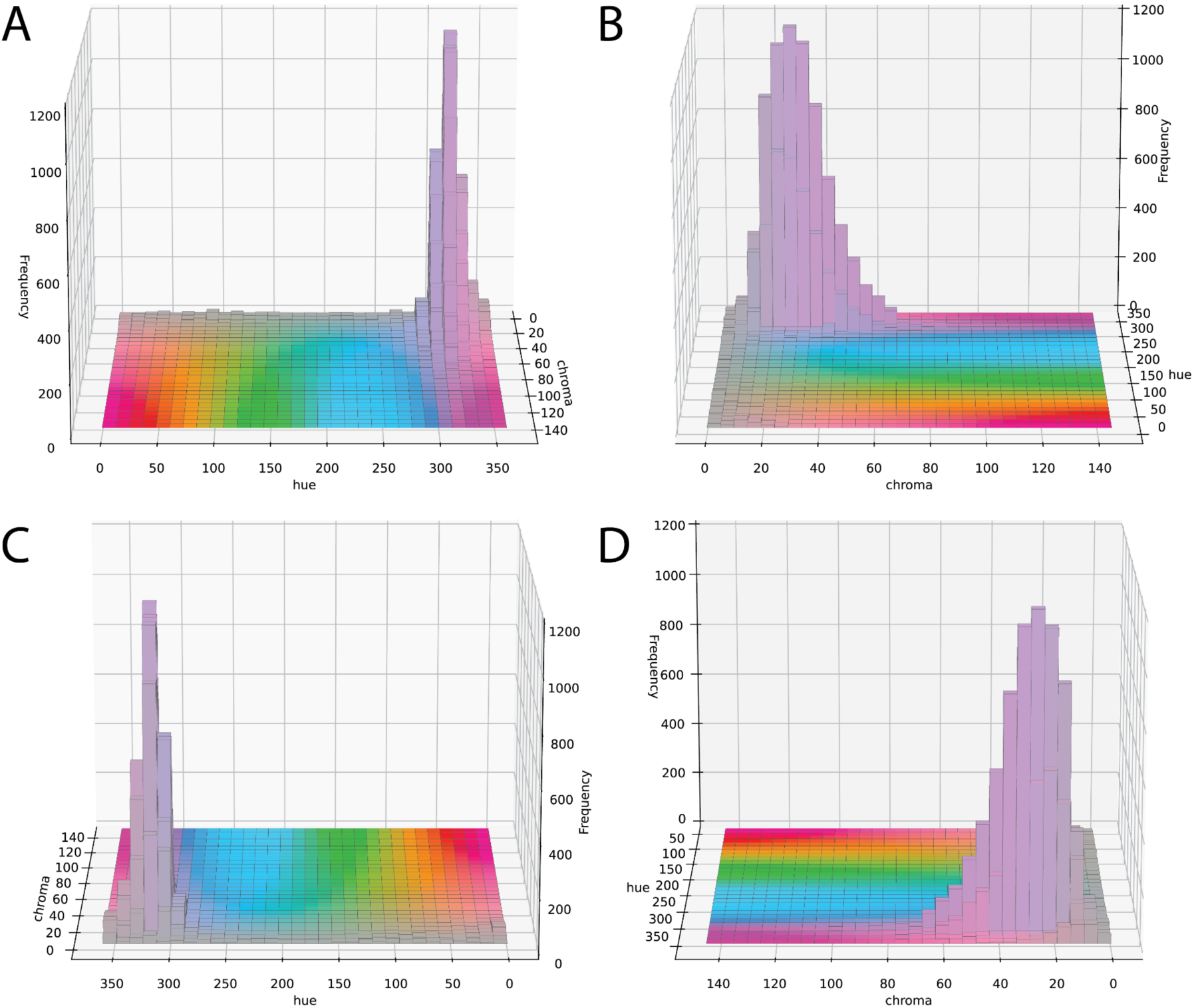
Complete rotation (A-D: 0°,90°,180°,270°) of the three-dimensional histogram from Figure 3A, showing frequency of observations across values of chroma and hue from LCh color space (lightness held constant at value 70).

**Figure S4:**
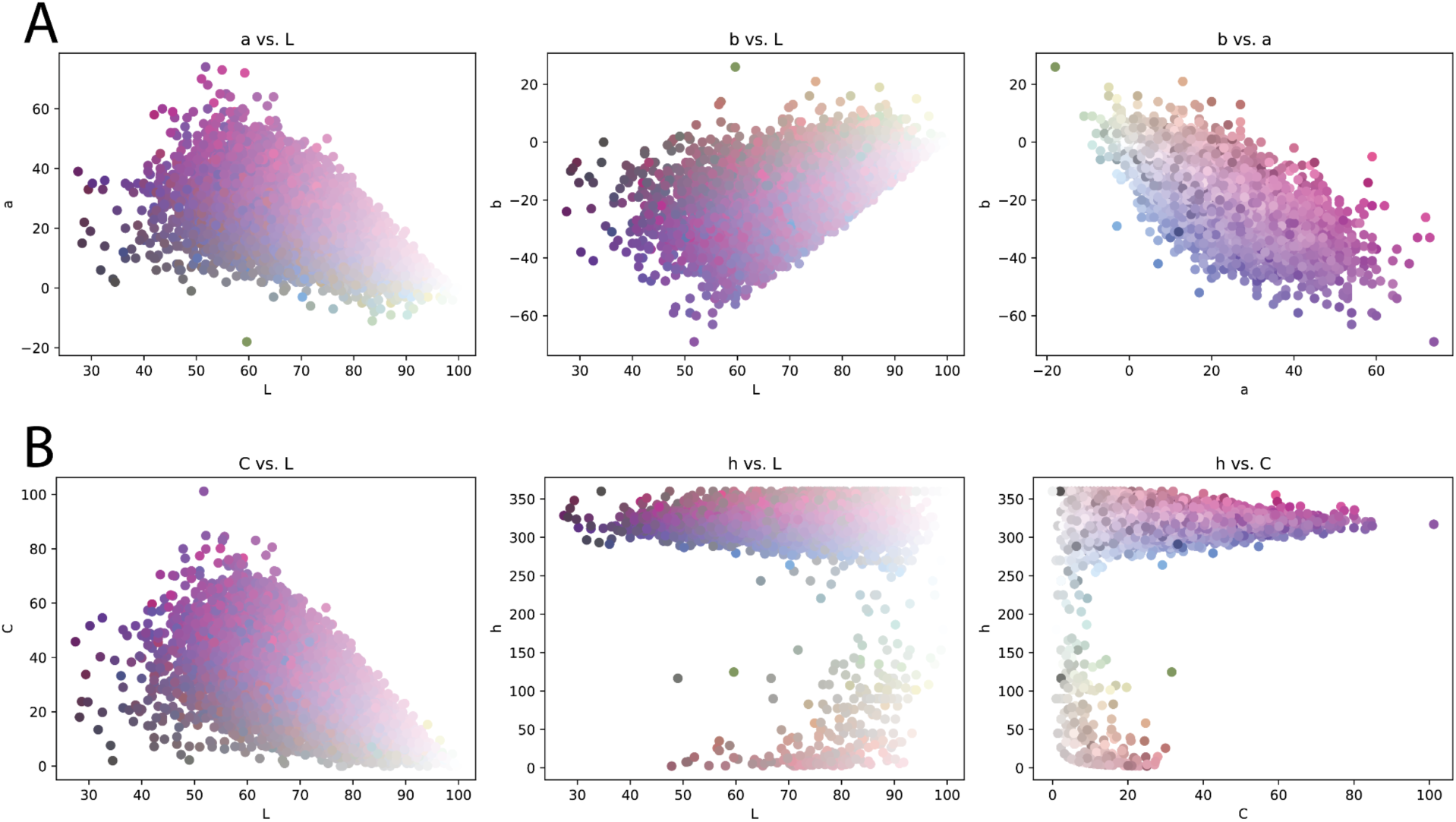
Pairwise projection of color components in A) CIELAB color space, and B) LCh color space. The bottom-right panel is identical to Figure 3B.

**Figure S5:**
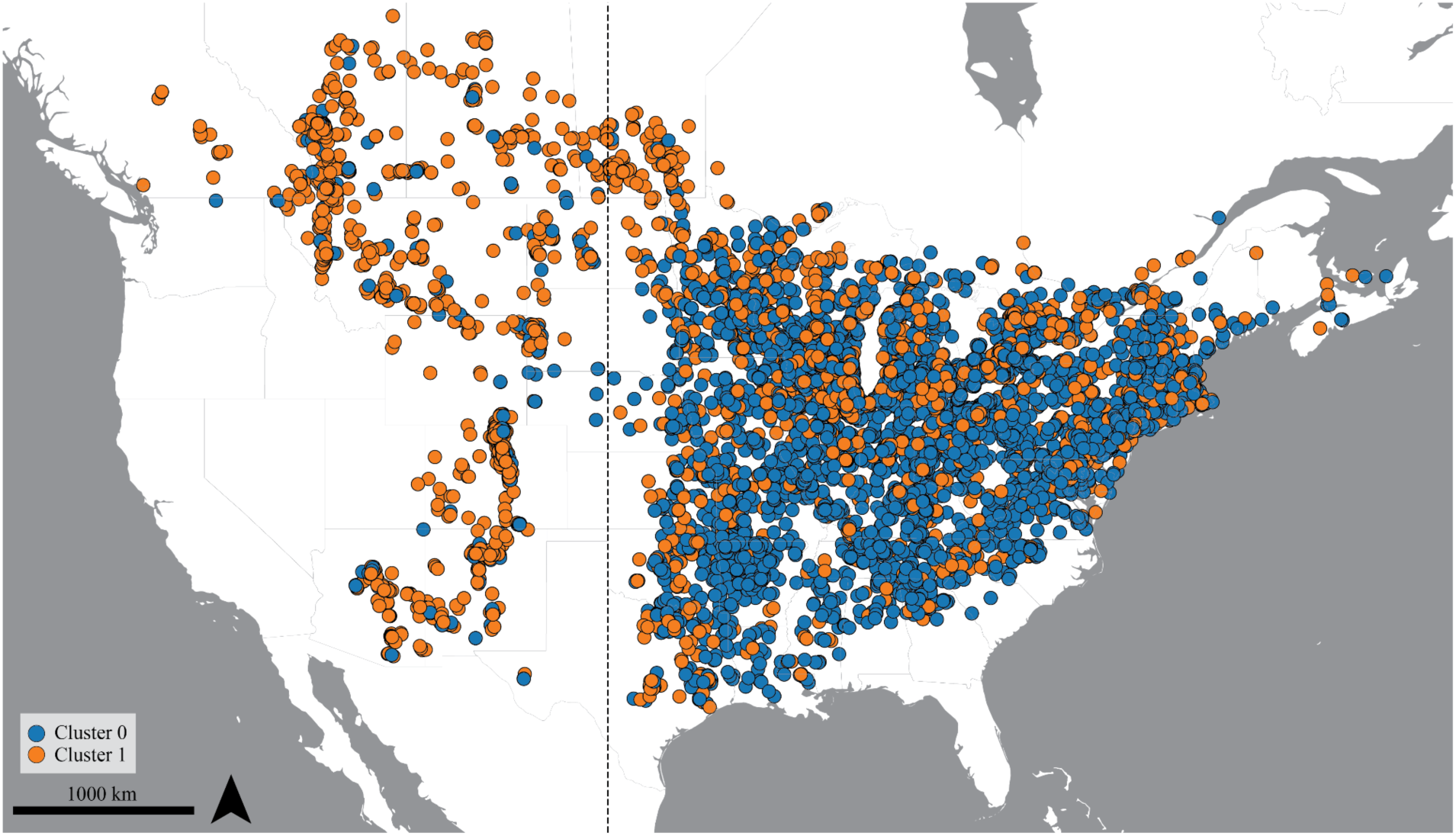
K-means clustering on the CIELAB color components with k=2. Color of point represents cluster identity for each observation. Sliding window averaging from this data across longitudes produced Figure 3D. The dotted line indicates -100° longitude.

